# RluA is the major mRNA pseudouridine synthase in *Escherichia coli*

**DOI:** 10.1101/2023.12.08.570740

**Authors:** Cassandra Schaening-Burgos, Gene-Wei Li, Wendy Gilbert

## Abstract

Pseudouridine (Ψ) is an ubiquitous RNA modification, present in the tRNAs and rRNAs of species across all domains of life. Conserved pseudouridine synthases modify the mRNAs of diverse eukaryotes, but the modification has yet to be identified in bacterial mRNAs. Here, we report the discovery of pseudouridines in mRNA from *E. coli*. By testing the mRNA modification capacity of all 11 known pseudouridine synthases, we identify RluA as the predominant mRNA-modifying enzyme. RluA, a known tRNA and 23S rRNA pseudouridine synthase, modifies at least 31 of the 44 high-confidence sites we identified in *E. coli* mRNAs. Using RNA structure probing data to inform secondary structures, we show that the target sites of RluA occur in a common sequence and structural motif comprised of a ΨURAA sequence located in the loop of a short hairpin. This recognition element is shared with previously identified target sites of RluA in tRNAs and rRNA. Overall, our work identifies pseudouridine in key mRNAs and suggests the capacity of Ψ to regulate the transcripts that contain it.

**Author Summary:** While RNAs are initially transcribed using only the nucleotides A, G, C and U, these can be enzymatically modified into many different post-transcriptional modifications. tRNAs and rRNAs across all domains of life are modified with pseudouridine, an isoform of uridine that is inserted by highly conserved enzymes. In many eukaryotes, it has been demonstrated that some of these enzymes can also insert pseudouridines in mRNA, where they are poised to impact gene expression through their effects on the transcript. Here we show that protein-coding transcripts in *E. coli* are also modified with pseudouridine, indicating that mRNA pseudouridylation is also a highly conserved activity. RluA is the main mRNA pseudouridine synthase, introducing the modification into the transcripts of dozens of protein coding genes with high specificity. Its target sites are defined by a combined sequence and secondary structure motif. Two additional enzymes, RluC and RluD, introduce a few additional sites. All three of these enzymes belong to the same sub-family of pseudouridine synthases, and homologs of these also modify mRNAs in humans. Thus, mRNA modification by these enzymes might be a conserved activity with the capacity to impact gene regulation.

## Introduction

All organisms modify their tRNAs and rRNAs extensively, introducing dozens of unique RNA modifications in many positions. It has become clear that mRNAs are also modified, as evidenced by the discovery of at least nine distinct modifications in the mRNAs of organisms across all domains of life (1–8, 46, 47). Pseudouridine (Ψ) in particular appears especially widespread, having been identified in the mRNAs of human, mice, yeast, *Trypanosoma brucei*, and *Arabidopsis thaliana* (1,2,9,10). However, it has yet to be detected in bacterial mRNAs.

Pseudouridine is an isomer of uridine, generated by rotating the uridine base to obtain an additional hydrogen bond donor and replacing the C-N glycosidic linkage with a C-C bond, while maintaining the same Watson-Crick face (Fig 1a) (11). Pseudouridylation is carried out by a universally conserved class of enzymes called pseudouridine synthases, which insert the modification in tRNAs and rRNAs, as well as other non-coding RNAs (12,13). In eukaryotes, some of the same enzymes that modify these canonical targets also insert pseudouridines into mRNAs (2,14,15). Because the pseudouridine synthases in *E. coli* belong to this conserved family, it is plausible that their mRNAs are also being modified with Ψ.

**Fig 1.**
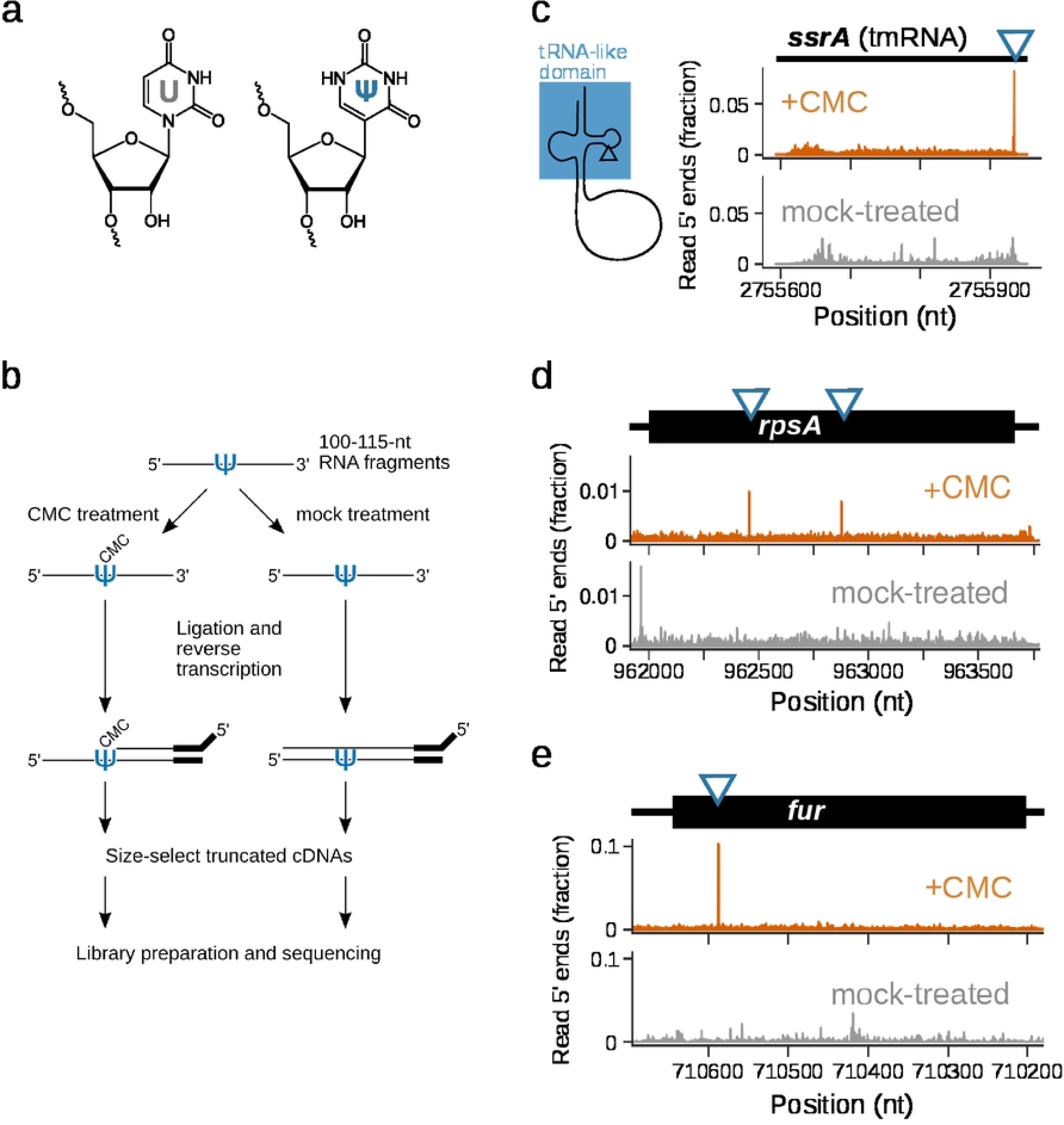
Pseudo-Seq identifies known and novel pseudouridine sites in the E. coli transcriptome. (a) Structures of uridine and pseudouridine. (b) Diagram of the PseudoSeq method. Treatment with CMC causes robust reverse transcription (RT) stops at pseudouridine sites. These are enriched via size selection and captured for sequencing. (c) Validation of a known site in the tmRNA gene, ssrA. Its tRNA-like domain contains a pseudouridine, indicated on the gene track and secondary structure diagram. Data tracks show read 5′ ends, normalized to total transcript reads, in wild-type CMC-treated samples (orange) and mock-treated samples (gray). (d) Pseudo-seq signal for two novel mRNA sites on the rpsA mRNA and for (e) a novel site in the fur mRNA

Through its biochemical effects, Ψ is poised to affect multiple stages in the life cycle of an mRNA. Pseudouridine has been consistently shown to stabilize RNA duplexes, including secondary structures and intermolecular RNA:RNA interactions (16–18). Through this mechanism, pseudouridine has the potential to modulate regulatory structures in mRNA, such as riboswitches, as well as interactions with small regulatory RNAs. Ψ can also impact the conformation of unpaired RNA regions by enhancing base stacking (19).

Pseudouridine can inhibit RNA-protein interactions by conferring additional backbone rigidity. In vitro, this has been demonstrated with pseudouridylated oligos and at least three different RNA-binding proteins, each of which had reduced affinity for pseudouridylated RNA compared to unmodified RNA (20–22). Similarly, modification can affect the activity of enzymes acting on RNA; recent work showed that the ribonuclease RNase E cleaves pseudouridylated RNAs less efficiently than unmodified ones, and did so at altered positions (23). Lastly, there is growing evidence for effects on translation fidelity: pseudouridine in stop codons can allow for stop-codon readthrough (24), whereas pseudouridine in some sense codons has been shown to enable translational recoding (25). This variety of effects highlights an important principle: the impact of pseudouridine on a given mRNA is likely to be highly context-dependent and subject to adaptive evolution, based on its interactome and local sequence and structure.

Here, we extensively characterize the pseudouridylation profile of exponential-phase *E. coli*, and determine the mRNA modification capacity of each of its 11 pseudouridine synthases. We discovered that mRNA pseudouridylation is almost exclusively carried out by the synthase RluA, posing it as a novel regulator of mRNA processing and function. This enzyme introduces pseudouridines in at least 31 mRNA sites, and is likely to modify additional sites that are lowly expressed. We characterize the sequence and structural context of the sites modified by RluA and determine that they occur in a common ΨURAA motif, often positioned within a small secondary structure element which resembles the ribosomal target site of RluA.

## Results

### Mapping pseudouridines by Pseudo-Seq

Pseudouridine detection is enabled by chemical treatment that generates a modification-dependent signature in sequencing datasets, even though Ψ by itself cannot be detected through reverse transcription and sequencing. One such reagent is CMC (N-cyclohexyl-N′-β-(4-methylmorpholinium)ethylcarbodiimide p-tosylate), a bulky carbodiimide that derivatizes the NH groups in Ψ and causes robust termination during reverse transcription (26). To detect pseudouridines in the *E. coli* transcriptome, we used the Pseudo-Seq method (1), which combines CMC treatment with size-selection steps that enrich for truncated reverse transcription products, followed by high-throughput sequencing (Fig 1b). This generates a CMC-dependent pileup of reads whose 5′ ends are immediately downstream of the modified site (Fig 1c).

We applied Pseudo-Seq to wild-type *E. coli* grown to exponential phase (OD600 ∼ 0.10–0.15). This approach successfully reproduced a known site in the tRNA-like domain of tmRNA (Fig 1c) (27), as well as several known sites in tRNAs and rRNAs (Fig S1). Each of these sites exhibits the expected CMC-dependent peak at the annotated position, indicating that known sites are reliably captured in our dataset.

### *E. coli* mRNAs are abundantly modified with pseudouridine

To identify pseudouridines in *E. coli* mRNAs along with the responsible pseudouridine synthases, we generated Pseudo-Seq datasets for wild-type *E. coli* and for individual knockout strains of all 11 pseudouridine synthases. In the wild-type dataset, we observed strong CMC-dependent peaks in some mRNAs, indicating the presence of pseudouridines in protein-coding genes (Fig 1d,e). In order to systematically call Ψ sites transcriptome-wide while accounting for biases arising from nearby modifications or transcript ends, we implemented a modified Z-score that includes 95% winsorization and background window adjustment (Fig S2, Methods). We identified candidate pseudouridine sites as strong peaks in the CMC-treated library that were absent from the mock-treated library. Based on the distribution of Z-scores among uridines in mRNAs, we defined these strong peaks as those that have a Z-score greater than 15 (Fig 2a) and at least 3-fold greater than the peak in the mock-treated library (Fig 2b). Using these stringent criteria, we were able to validate 9 out of the 10 ribosomal RNA sites, with no false-positives (Fig S2c).

**Fig 2.**
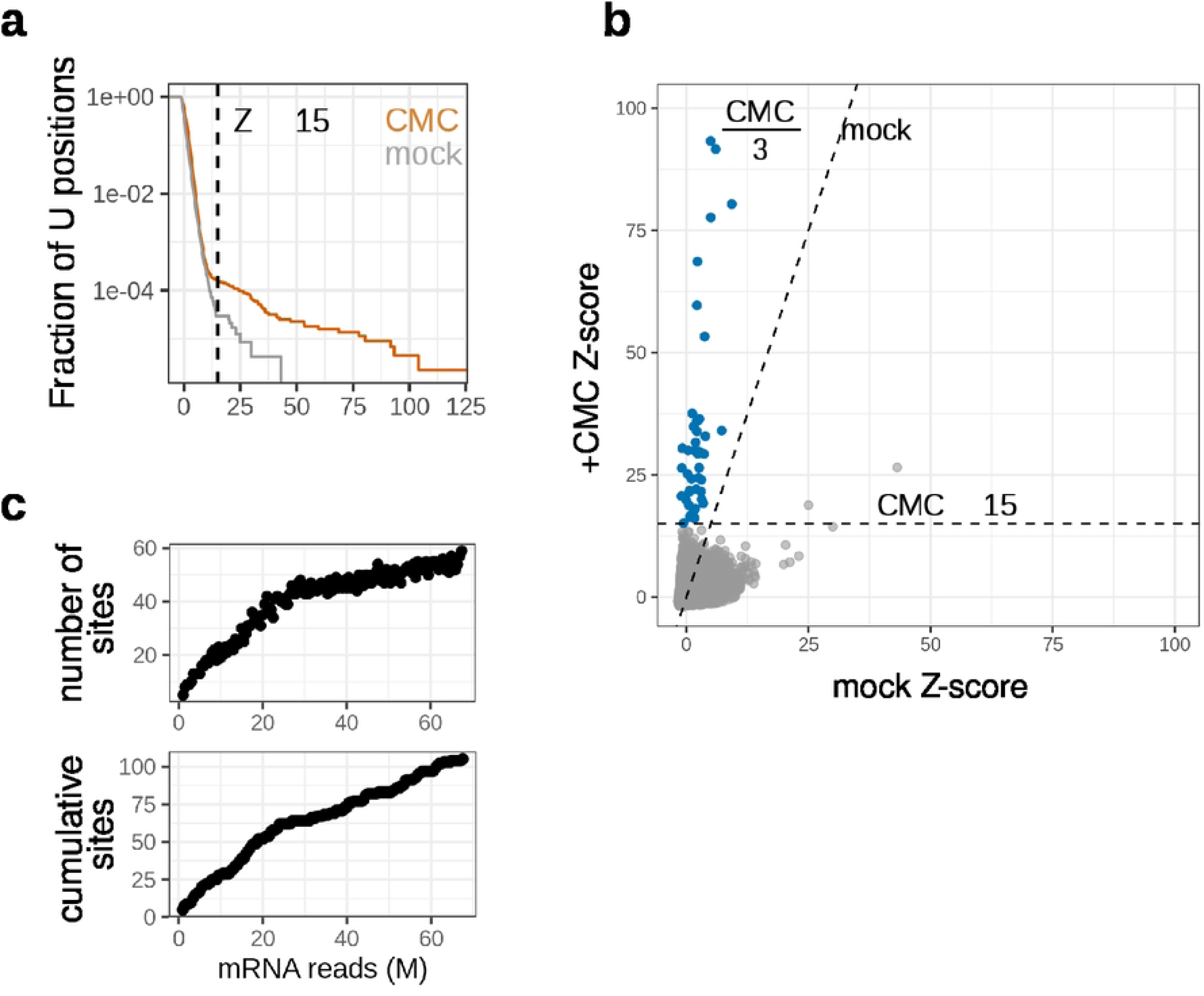
Systematically calling pseudouridine sites transcriptome-wide. (a) Z-score distribution for all uridines in mRNA in wild-type libraries, with CMC-treated in orange and mock-treated in gray. Distribution is visualized as an 1 − cumulative density function, indicating the fraction of positions that have a Z-score greater than a given value. The cutoff for calling a peak in the CMC-treated library, Z ≥ 15, is indicated by the dashed line. (b) Joint distribution of mRNA Z-scores in CMC-treated (y-axis) and mock-treated (x-axis) libraries. Cutoffs for calling potential pseudouridine sites are indicated by the dashed lines: sites must have a Z-score ≥ 15 in the CMC-treated library, and must be at least 3-fold greater than the corresponding peak in the mock-treated library. Sites that meet both cutoffs are considered potential pseudouridine sites and are shown in blue. (c) Number of CMC-dependent peaks called at increasing mRNA read coverage. Top panel shows the number of peaks called in each subsample, bottom panel shows the cumulative number of unique peaks.

We then applied the modified Z-score metric to reads mapped to the entire *E. coli* transcriptome (Fig 2b, Methods). Peak height calculations are very noisy at lower sequencing coverage (Fig S2d), necessitating strict coverage cutoffs in the background region surrounding each putative site. Applying these cutoffs limits site calling to genes with sufficient expression, and so the number of sites called in a library will vary depending on sequencing depth. To address this issue, we have generated a high-depth wild-type Pseudo-seq library, which spans 24.2% of uridines in protein-coding genes, to call a high-confidence list of sites. The profiled uridines encompass 1532 protein-coding genes, out of the 4099 detected in our sequencing data. The genes sequenced to sufficient depth for pseudouridine detection were expressed to a median RPKM of 201, while the overall median expression level was 44.4 (Fig S2E).

This top-tier list of targets consisted of 44 mRNA sites (Fig 2c). 43 of these were within coding-sequences, where they could potentially impact the fidelity and rate of translation. Modifications outside of the coding sequence were comparatively rare, as expected from the relative sizes of these regions in *E. coli*, with only one modification in the 5′ UTR, and none in the 3′ UTR.

The genes modified with pseudouridine do not appear to form a coherent regulon—rather than targeting genes that are part of the same processes or pathways, the modification is present in mRNAs encoding components of many different biological processes. Among these, there is modification of genes that encode proteins involved in translation, including three ribosomal proteins (RpsA, RpsH, RpsQ), two translation elongation factors (TufA, FusA), and one tRNA biogenesis factor (Fmt) (Supplementary Table 1).

Sequence coverage is the limiting factor in calling new sites. Thus, most called sites are in highly expressed genes because these were likeliest to have sufficient coverage. To determine whether pseudouridylation also occurs in mRNAs with lower expression levels, we simulate higher coverage by pooling reads from different Pseudo-Seq libraries. Increasing coverage in this manner identified many new CMC-dependent peaks, with up to 100 sites detected when coverage exceeds 65 million mRNA-mapping reads, respectively, indicating that pseudouridine is also present in genes with lower expression levels. (Fig 2c, Supplementary Table 2).

### RluA is the predominant mRNA pseudouridylation enzyme

Pseudouridine is installed in *E. coli* tRNAs and rRNAs by a universally conserved class of pseudouridine synthases. These enzymes are derived from an ancestral pseudouridine synthase, identified by a common fold, and belong to five different families (with an additional sixth family in eukaryotes) (13,28). Each of these five families is represented among the *E. coli* pseudouridine synthases, with two of these families having diversified further, for a total of 11 pseudouridine synthases. To determine which of these enzymes carry out mRNA pseudouridylation, we obtained knockout strains of each of these synthases and carried out Pseudo-Seq on cultures grown to exponential phase (OD600 ∼ 0.10–0.15).

Enzyme-dependent sites are expected to show a CMC-dependent peak across all libraries except the relevant knockout (Fig 3a,b, Methods). In this manner, 31 out of the 44 high-confidence sites are unambiguously assigned to RluA. There is additional modification carried out by the enzymes RluC and RluD (Fig 3c); RluD unambiguously modifies at least three mRNA sites, while RluC modifies a single site in the *atpG* mRNA. Each of these three enzymes is known to modify multiple sites among tRNAs and rRNAs. RluA modifies four tRNAs and a single position in rRNA; the RluD enzyme was previously known to modify three sites in the 23S rRNA, all in the loop of helix 69; and RluC modifies three sites in separate regions of the 23S rRNA. Meanwhile, the remaining synthases appear to be highly specific for a single rRNA site, or to a specific position in tRNAs.

**Fig 3.**
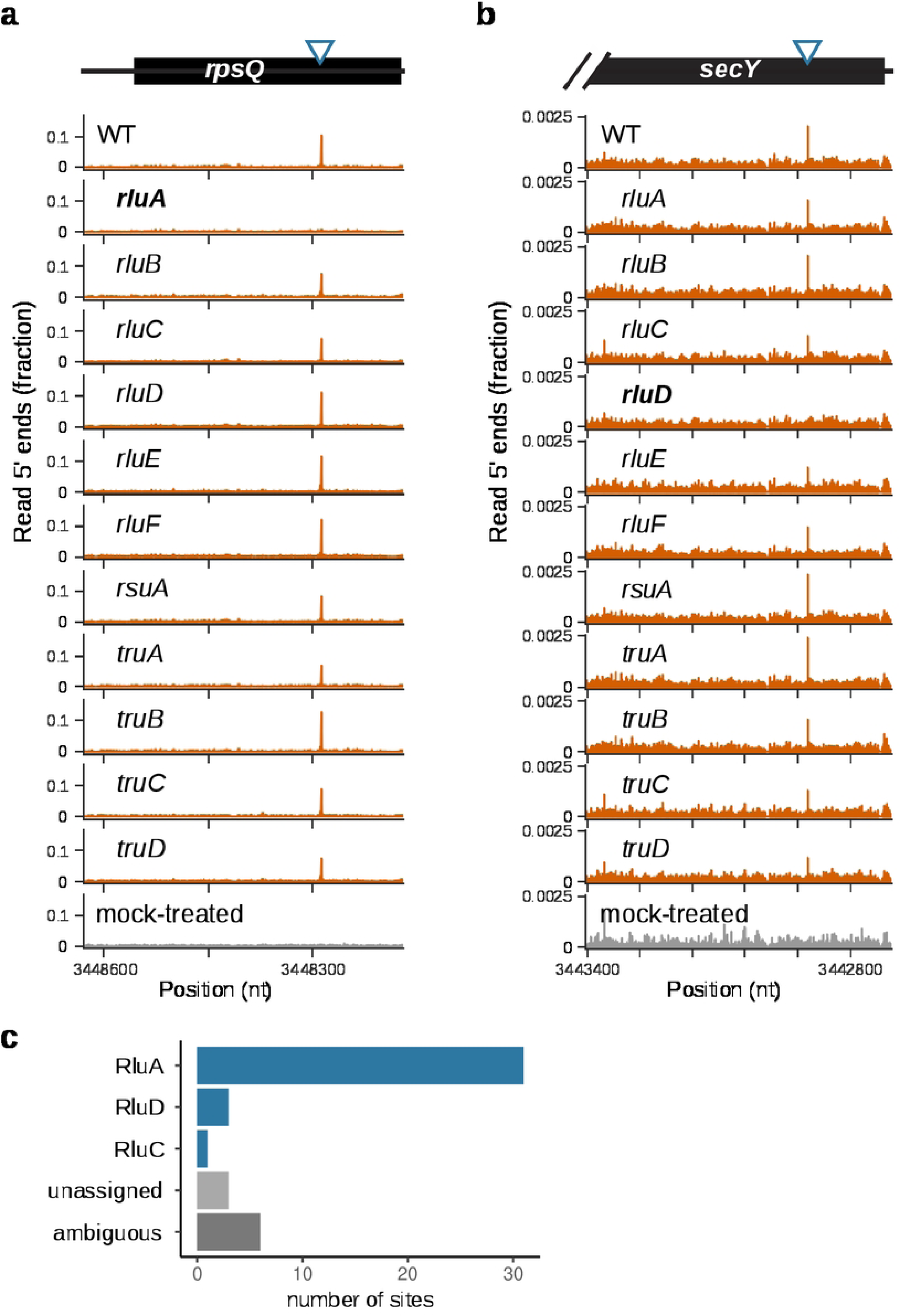
RluA is the predominant mRNA pseudouridine synthase in E. coli. (a) Pseudo-Seq signal for a sample mRNA site in 13 libraries: wild-type, all 11 pseudouridine synthase knockouts, and a mock-treated control for the wild-type. The site appears as an RluA- and CMC-dependent peak in read 5′ ends. (b) Pseudo-Seq signal for a sample RluD-dependent site in an mRNA. The site appears as a RluD- and CMC-dependent peak in read 5’ ends. (c) Enzyme assignment for the highest confidence sites. In order to assign enzymes, we required sufficient coverage in at least 7 of the 12 CMC-treated libraries, and for the peak to be significantly dependent on CMC.

Nine of the high-confidence pseudouridine sites could not be conclusively assigned to an enzyme. In these cases, low coverage of the target site in a subset of the libraries prevented unambiguous assignment. Sites remain unassigned (Methods) when they exhibit a strong peak in all libraries where they have high enough coverage to call a CMC-dependent peak in the wild-type library, but insufficient coverage in one or more knockouts prevents the absence of a peak from being confidently called. Meanwhile, sites can be ambiguously assigned when the peak is absent in two or more knockout libraries. This could be biological, a consequence of partially redundant modification by more than one enzyme, or technical, resulting from low coverage.

We performed RNA sequencing on wild-type and Δ*rluA* to examine the effects of RluA on expression of its mRNA targets. We identified few changes overall. A single mRNA target of RluA, *fimA/*the *fim* operon, was significantly affected. The 5’ UTR of the *fim* operon contains an RluA-dependent pseudouridine site (Fig 4a). The *fim* operon encodes a polycistronic transcript with the components of the Type 1 pilus, which are rod-like structures on the outer membrane of *E. coli* that promote surface adhesion. *fimA*, which encodes the major subunit of the pilus, experienced the greatest fold-change in expression, though all genes in the operon were upregulated. This upregulation was highly specific for the *fim* operon, as the *rluA* knockout did not result in global changes in gene expression (Fig 4a). One additional gene, *pdeL*, was substantially upregulated despite not being a modification target of RluA.

**Fig 4.**
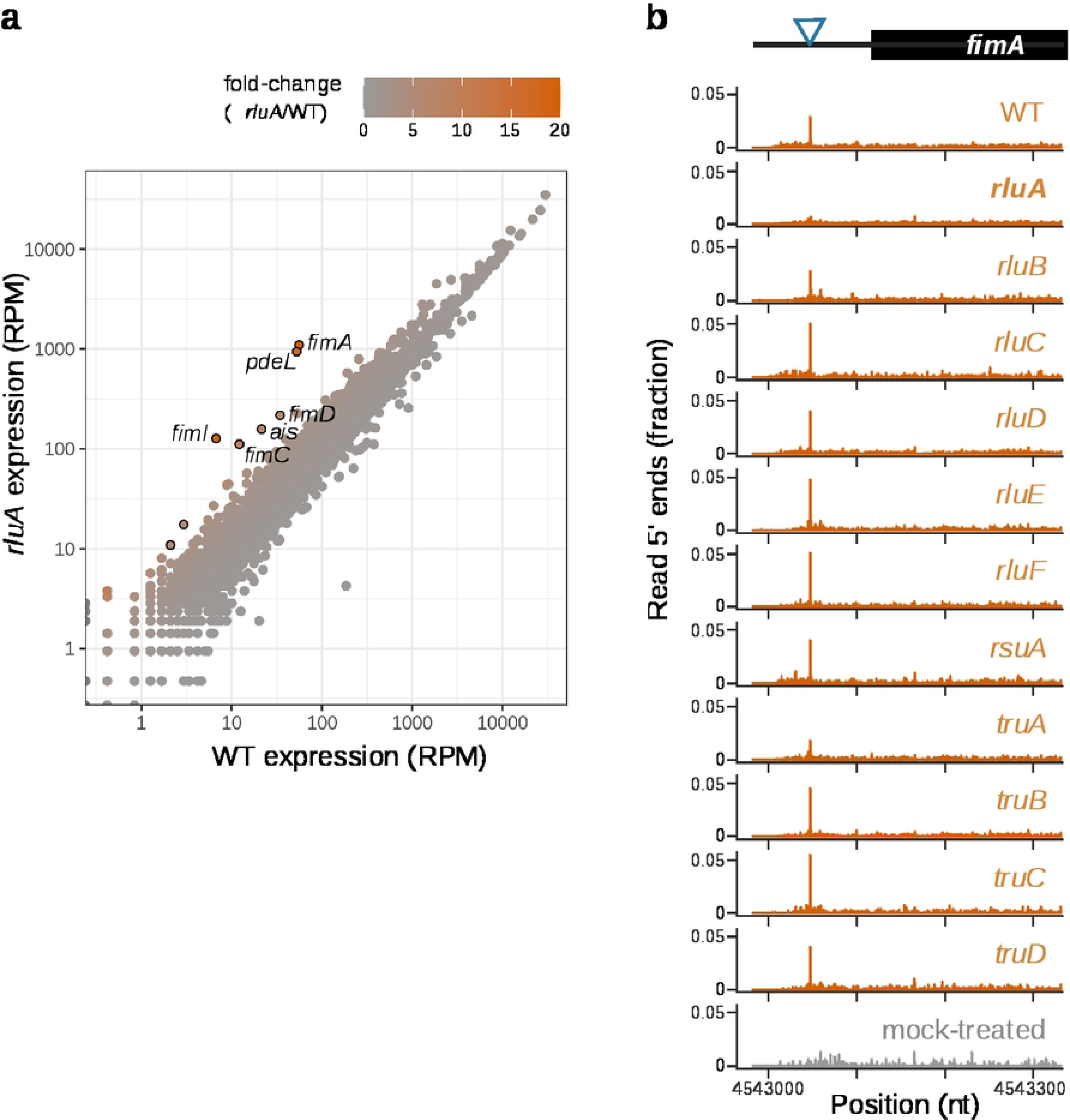
Loss of RluA alters gene expression. (a) Gene expression data in wild-type (x-axis) and ΔrluA (y-axis). Genes are colored by their fold change in expression relative to wild-type. RPM: reads per million. (b) Pseudo-Seq read 5′ end counts for CMC treated libraries (orange) in every pseudouridine synthase knockout background, and in a mock treated library (gray). Counts are expressed as a fraction of reads mapped to the transcript.

### RluA modifies positions in mRNA, tRNA, and rRNA that share a sequence and structure motif

Our results illustrate the diversity of pseudouridine synthase targets, which span multiple classes of RNA. Strikingly, the enzyme RluA is capable of specifically modifying sites in tRNA, rRNA and mRNA, despite the drastic differences in structure and life cycle among these classes. By contrast, most of the other *E. coli* pseudouridine synthases modify sites in a single region or position. We next show that RluA modification across these distinct sites are guided by a common sequence and structure motif.

RluA’s canonical targets, which consist of one site in the large subunit rRNA and position 32 in the anticodon stem-loop for four different tRNAs (31,32), occur within a ΨURAA motif (where R indicates an A or a G), in which positions U1 and A4 are particularly important for recognition (Fig 5a)(32). We observe that the high-confidence targets of RluA occur in a similar context as the canonical sites, with highly constrained U1 and A4 positions, and a strong preference for A at +3 and +5 (Fig 5b). This sequence motif is also enriched among the ambiguous and unassigned sites sites (Fig 5b), indicating that RluA is likely to modify more sites than those we have identified under our stringent hit-calling criteria.

**Fig 5.**
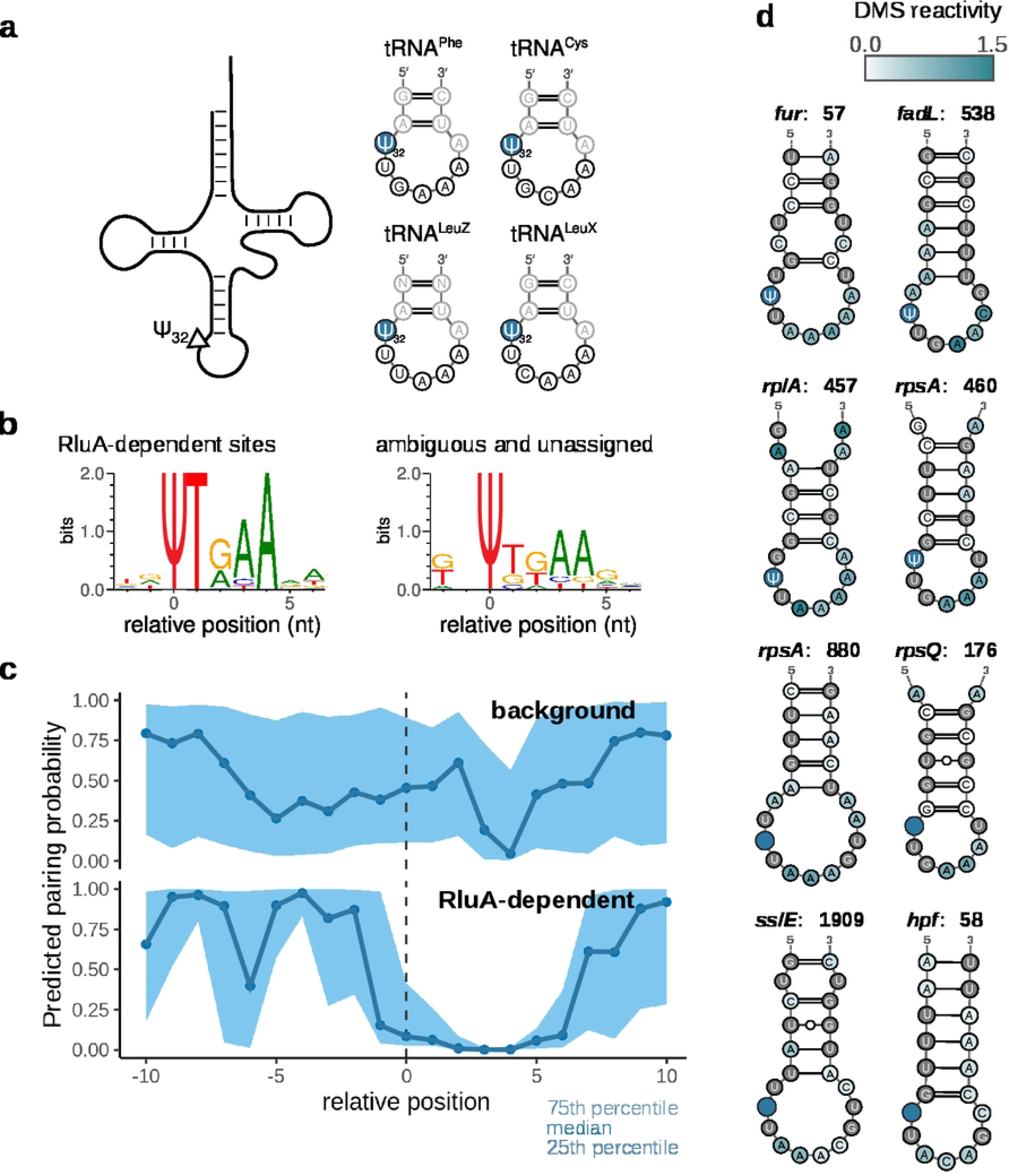
RluA targets tRNAs, rRNA, and mRNAs with a common sequence and structure motif. (a) Sequence and structure of all four canonical tRNA targets of RluA. (b) Sequence logos illustrating the sequence motif for the high confidence RluA-dependent mRNA sites (left, N=31) and for ambiguous and unassigned sites (right, N=9) (c) Predicted pairing probabilities for the mRNA targets of RluA. DMS-seq reactivities in the 100-nt surrounding each site were obtained from Burkhardt et al (2017) and used to inform secondary structure prediction. Background set consists of 94 highly expressed sites containing the UURAA motif, but lacking a CMC-dependent peak. (d) Sample structures of RluA target mRNAs. Shades of green indicate DMS reactivity, and the pseudouridine site is highlighted in blue.

The UURAA motif is ubiquitous in the transcriptome, with over 1898 instances sequenced to sufficient depth in this dataset, but the majority of these instances did not exhibit a CMC-dependent peak (Fig S4). Therefore, this simple motif cannot be sufficient for modification and additional features beyond the presence of a sequence motif must be required for recognition. The tRNA and rRNA targets of RluA share a similar structure, with the pseudouridines located in the loops of short hairpins, within one or two nucleotides of the last 5′-paired base (32).

Therefore, we sought to determine whether the mRNA targets of RluA occur in a similar structure context as the canonical sites in tRNA and rRNA. To determine the structural context of mRNA sites, we carried out dimethyl sulfate (DMS)-informed secondary structure prediction, using published RNA structure probing data from (33). These data provide DMS reactivity scores for each A and C nucleotide, with higher reactivity indicating more accessible nucleotides. Visualizing the distribution of predicted pairing probabilities for each position relative to the Ψ, we observe that the RluA mRNA sites follow a very similar pattern as the canonical tRNA and rRNA sites, suggesting a common structural motif (Fig 5c). Inspection of the predicted structures confirms that this is the case (Fig 5d, Fig S5).

## Discussion

Our work demonstrates that many important *E. coli* mRNAs are modified with pseudouridine, and that these modifications are overwhelmingly carried out by the synthase RluA. Under exponential growth in rich media, we identify 44 high-confidence sites in 42 highly-expressed genes, and estimate that there at least 80 total sites in mRNAs under these growth conditions. Because PseudoSeq relies on reverse transcription stops and ligation to capture pseudouridine sites, our results are not quantitative, and therefore we cannot infer what percentage of Us at a given position are pseudouridine. The extent of pseudouridylation at given positions may be quantified through targeted mutational mapping approaches (49, 50) or through direct RNA sequencing (48).

Prior to this study, very few modifications had been profiled in *E. coli* mRNAs, and only two types of modification had been conclusively located. Adenosine-to-inosine editing has been found in 15 transcripts, where they enable recoding of tyrosine codons. Meanwhile, N6- methyladenosine was found to be abundant in *E. coli* mRNAs, but the methyltransferases that insert them have not been identified, and its functions have not been elucidated. Other modifications known to occur in eukaryotic mRNAs have either not been profiled in *E. coli* (as is the case for N1-methyladenosine, dihydrouridine, and N4-acetylcytidine) or failed to be detected (5-methylcytosine).

Pseudouridine is almost exclusively introduced into *E. coli* mRNAs by RluA, with the exception of a few sites modified by RluD and RluC. RluA’s strong requirement for a UURNA motif, which is most often instantiated as UURAA, has deep implications for the distribution of pseudouridines in mRNAs. As a result of this sequence context, pseudouridines preferentially occur in valine (27% of high-confidence sites), leucine (24%), phenylalanine (20%), and isoleucine (13%) codons. In addition, the modification very rarely occurs in stop codons, with a single high-confidence event occurring in a stop codon. Meanwhile, very little can be elucidated about the binding requirements for the synthases RluC and RluD from this dataset due to the small number of total sites. A single RluC-dependent mRNA site was found, which shares a GNΨG sequence motif with the three RluC-dependent rRNA sites (Fig S3c). The three RluD- dependent sites mRNA sites are each directly preceded by a C, which is true of only one of its rRNA sites (Fig S3b). A previous study showed that purified RluD and RluC each modify many more sites *in vitro* than they do in cells (51), which suggests the participation of additional factors.

When binding its targets, RluA recognizes the target sequences by refolding them into a novel structure (32). A crystal structure of RluA in complex with the anticodon stem-loop of one of its target tRNAs shows that the bound sequence is rearranged to flip out bases A+2 and N+5 (positions relative to the pseudouridine) and the target uridine (U32 in tRNA) while inducing positions U+1 and A+4 to form a reverse-Hoogsteen base pair (32). These observations are consistent with the patterns we observe among mRNA targets: positions U+1 and A+4 are highly conserved among our sites, and the observed preference for the pseudouridines to occur in loops might facilitate assuming this induced conformation.

Our results indicate that RluA potentially binds to the mRNA transcripts of dozens of genes, which may by itself have consequences for the target RNA in addition to the direct properties of pseudouridine. Some RNA-modifying enzymes function as RNA chaperones (34,35), unfolding the target RNA and providing the opportunity to re-fold into the correct conformation. This activity has been demonstrated for the pseudouridine synthase TruB in its interactions with tRNA. Loss of TruB results in misfolding of tRNAs and a defect in aminoacylation. However, expressing a catalytic null mutant of TruB, able to bind to its tRNA targets but unable to modify them, restores normal tRNA function (34). It is possible that such a chaperone function may take place during mRNA modification by RluA, particularly if the modification occurs within a structured region, as we have demonstrated through analyzing RNA structure probing data.

We identified pseudouridylation of the 5′ UTR of *fimA* by RluA, as well as RluA-dependent expression in the *fim* operon. The *fim* operon encodes the components of the type 1 pilus (36). This operon is highly regulated, combining regulation of its promoter, control of two different isoforms and extensive secondary structure of its 5′ UTR. At the transcriptional level, expression of the operon can be turned on or off by recombinases that flip the upstream promoter, generating heterogeneity at this locus (37,38). In pathogenic strains of *E. coli*, regulation of this operon is key for virulence and adhesion. Interestingly, expression of RluA is significantly upregulated during infection of bladder epithelial cells (39). As this suggests a potential role for Ψ in virulence, uropathogenic *E. coli* strains may be a particularly interesting model system in which to explore the functions of mRNA pseudouridylation.

Loss of RluA could affect expression of the *fim* operon through various mechanisms: (a) lack of pseudouridylation could alter the structure and decay rate of the mRNA, (b) binding by RluA could carry out a chaperone function on the 5′ UTR, independently of modification, or (c) downstream effects from RluA could perturb regulation of this locus at the transcriptional level. Notably, one gene outside of the *fim* operon was differentially expressed to a similar extent as *fimA* (Fig 4a). This gene, *pdeL*, encodes a cyclic-di-GMP phosphodiesterase and transcriptional regulator. Until recently, it was only known to regulate its own transcription, but now additional targets have been identified by ChIP (40). Therefore, it will be important to determine whether *pdeL* carries out any regulation upstream or downstream of *fimA*, and whether pseudouridine plays a role in a regulatory network involving these genes.

Our results provide the first evidence for pseudouridine in bacterial mRNA. To identify phenotypes associated with these sites, it will be useful to survey growth and media conditions where loss of RluA confers a competitive disadvantage or growth defect, or where RluA is significantly differentially expressed. As with the example of uropathogenic *E. coli* infection, existing expression data from other virulent strains of *E. coli* may provide helpful leads. mRNA pseudouridylation by RluA family enzymes appears to be a highly conserved activity, as homologs RPUSD2 and RPUSD4 carry out modification of human pre-mRNAs (52).

## Figure Captions

**Fig S1.**
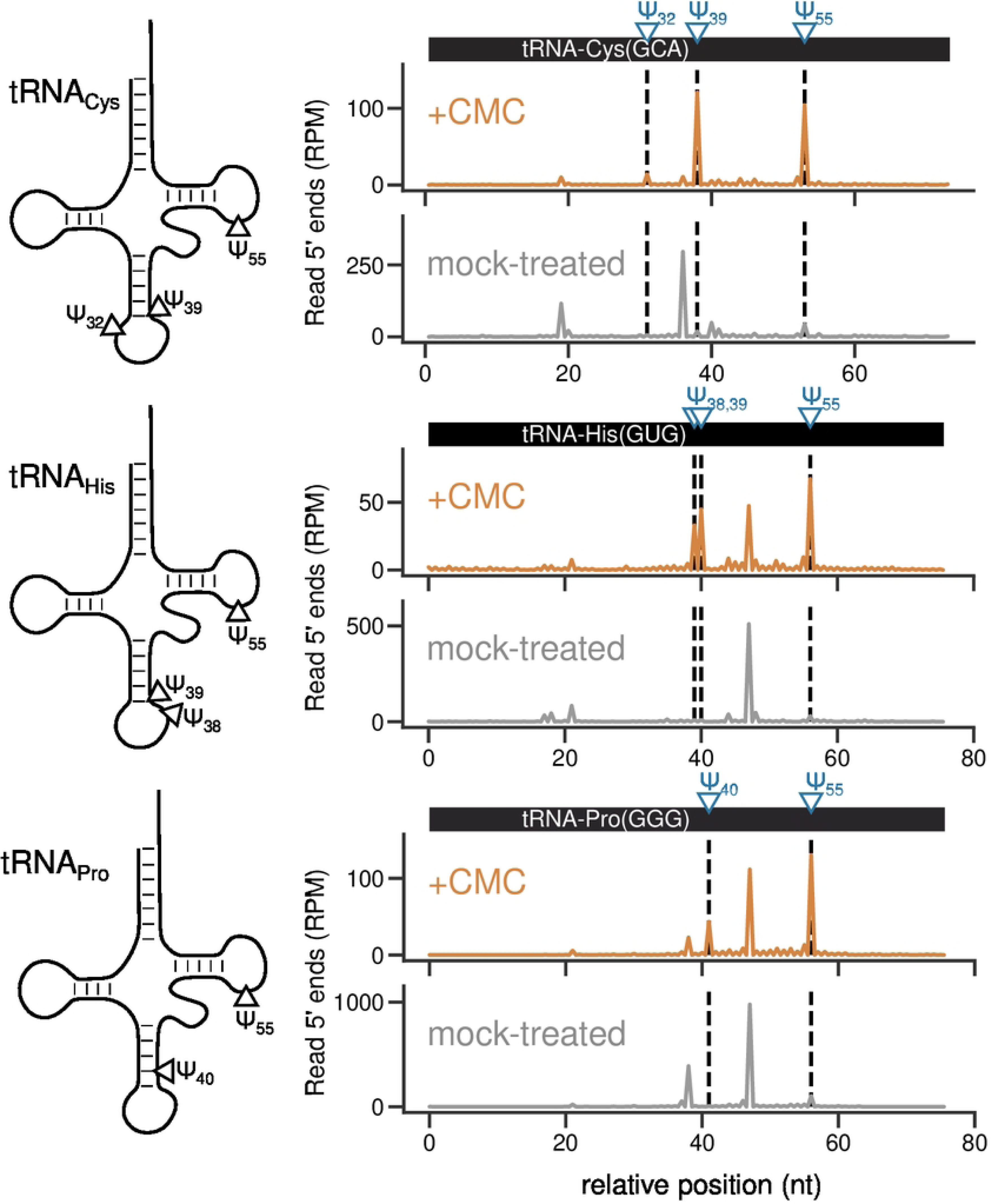
Validation of known sites in three tRNAs: tRNA-Cys(GCA) (top), tRNA-His(GUG) (middle), and tRNA-Pro(GGG) (bottom). PseudoSeq data tracks for CMC-treated (black) and mock-treated (gray) samples. Pseudouridine position is indicated with triangles on the secondary structure, and with blue dashed lines on the data track.

**Fig S2.**
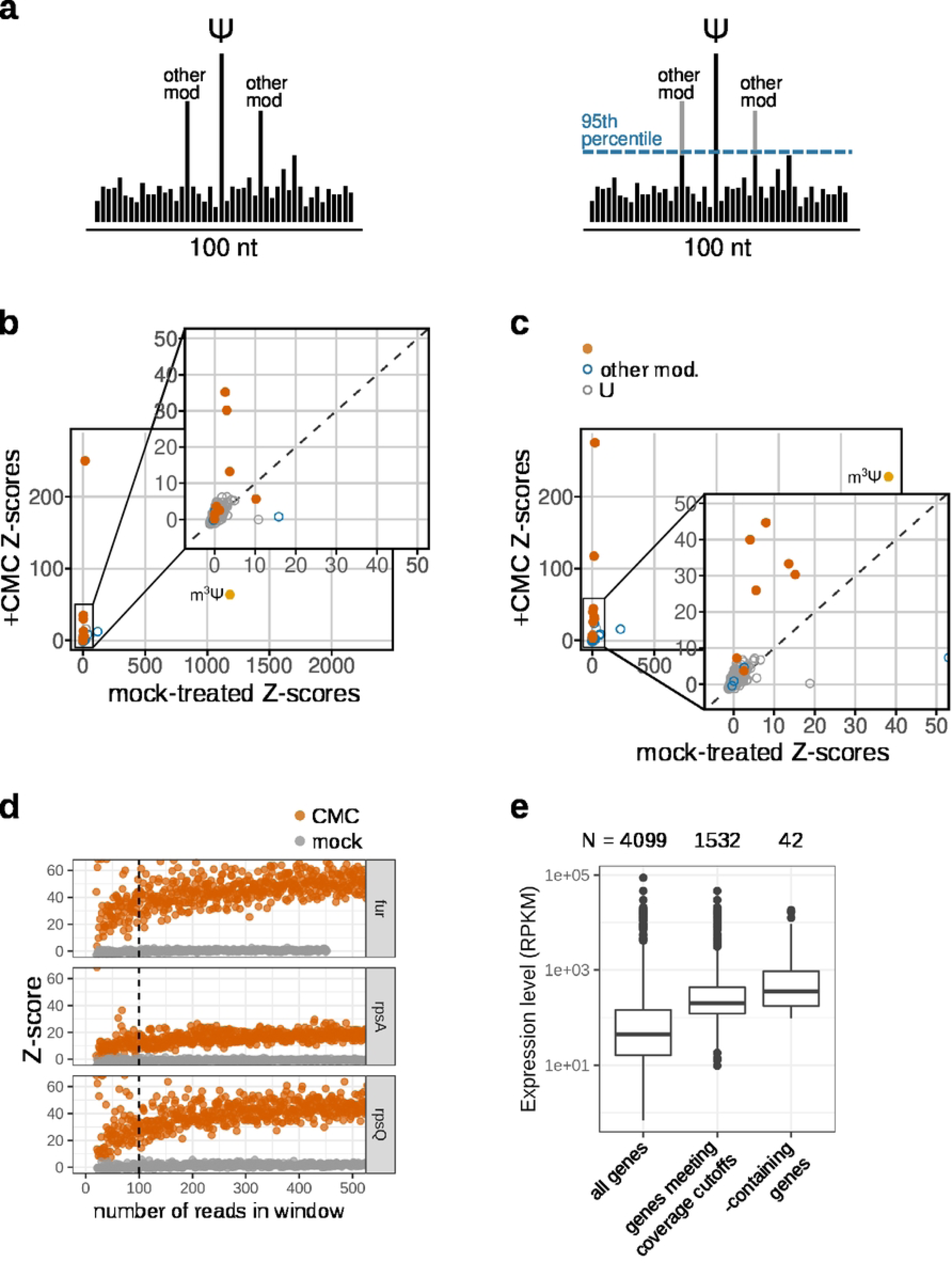
Z-score calculation before and after winsorization. (a) rRNA Z-scores calculated without background winsorization and with background winsorization. For (b) and (c) Z-scores for reference U-sites are shown and compared between a CMC-treated library and a mock-treated library. Known pseudouridine sites are colored in orange, other modified uridines are in blue, and unmodified uridines are in gray. (d) Selection of coverage cutoff. At high confidence sites, reads from a CMC-treated library (orange) and a mock-treated library (gray) were subsampled to various levels (x-axis shows the total number of reads mapping to the window after subsampling) and the corresponding Z-score was calculated (y-axis). Coverage cutoff of ≥ 100 reads is shown as dashed lines. (e) Expression level distributions for all expressed genes, genes that met coverage cutoffs in the wild-type pseudo-seq library, and genes containing a pseudouridine site. Gene counts are annotated at the top.

**Fig S3.**
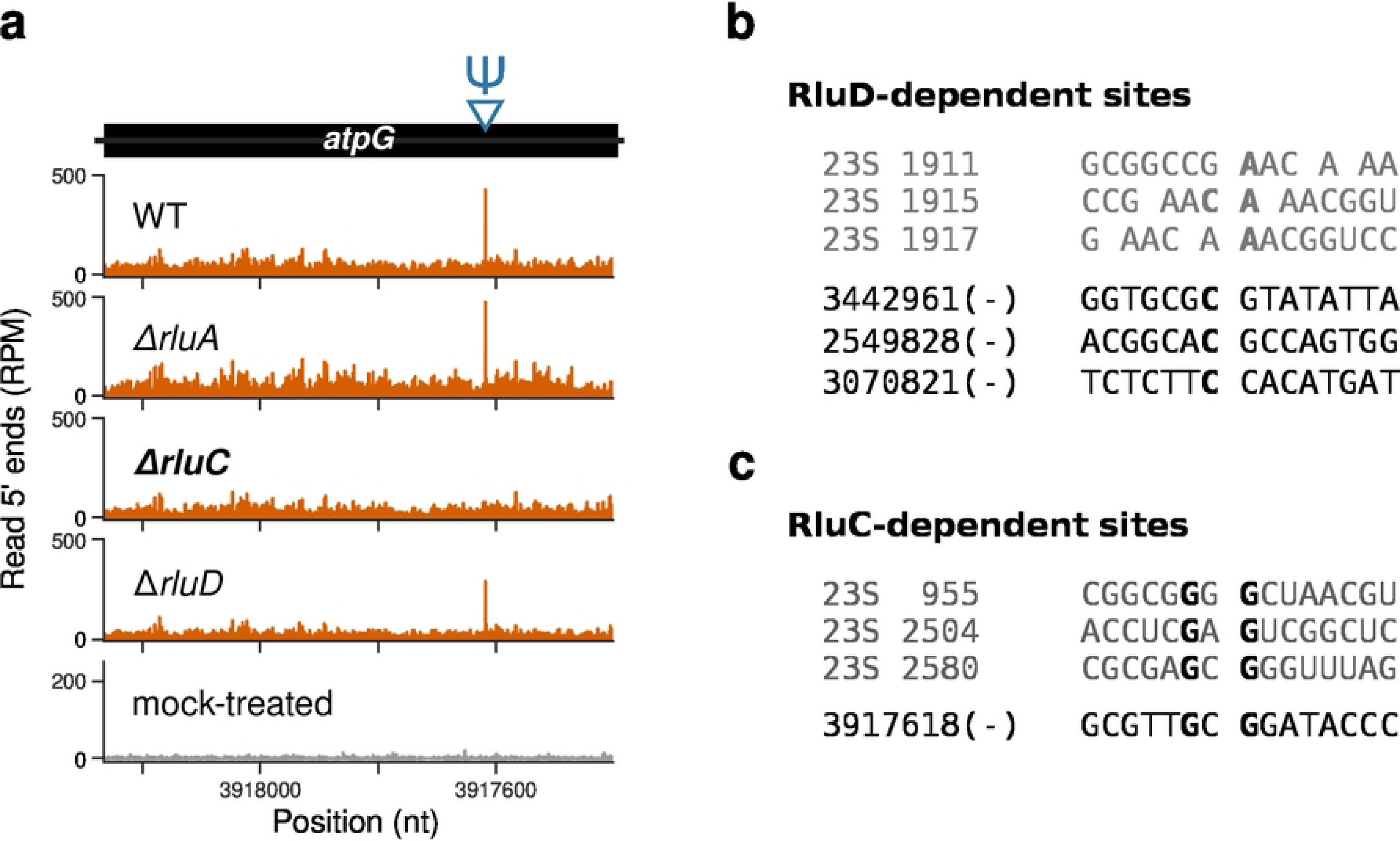
Reproducible RluC-dependent site in the atpG mRNA. (a) PseudoSeq signal for CMC-treated wild-type and CMC-treated knockouts of RluA, RluC, and RluD (orange) and for mock-treated (gray). The site appears as a RluC- and CMC-dependent peak in read 5’ ends. (b) Sequence alignment of RluD-dependent rRNA (gray) and mRNA (black) sites. (c) Sequence alignment of RluC-dependent rRNA and mRNA sites, with common nucleotides in bold

**Fig S4.**
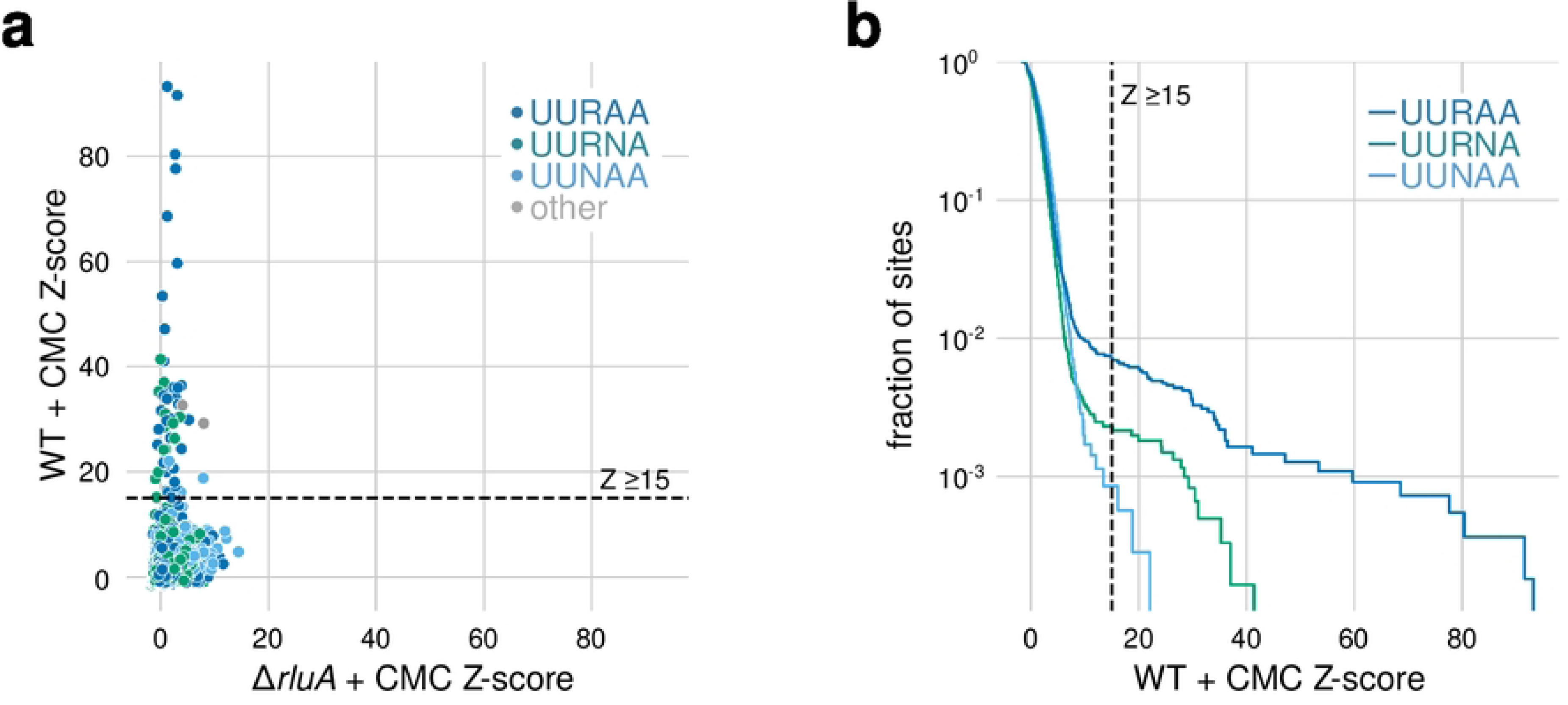
The UURNA motif is necessary but not sufficient for RluA-dependent pseudouridylation. (a) Pseudo-Seq signal (Z-score) for UURNA instances in CMC-treated wild-type (y-axis) vs ΔrluA (x-axis) samples. (b) Z-score distribution for UURAA, UUR[G|C|U]A, and UUYAA motif instances.

**Fig S5.**
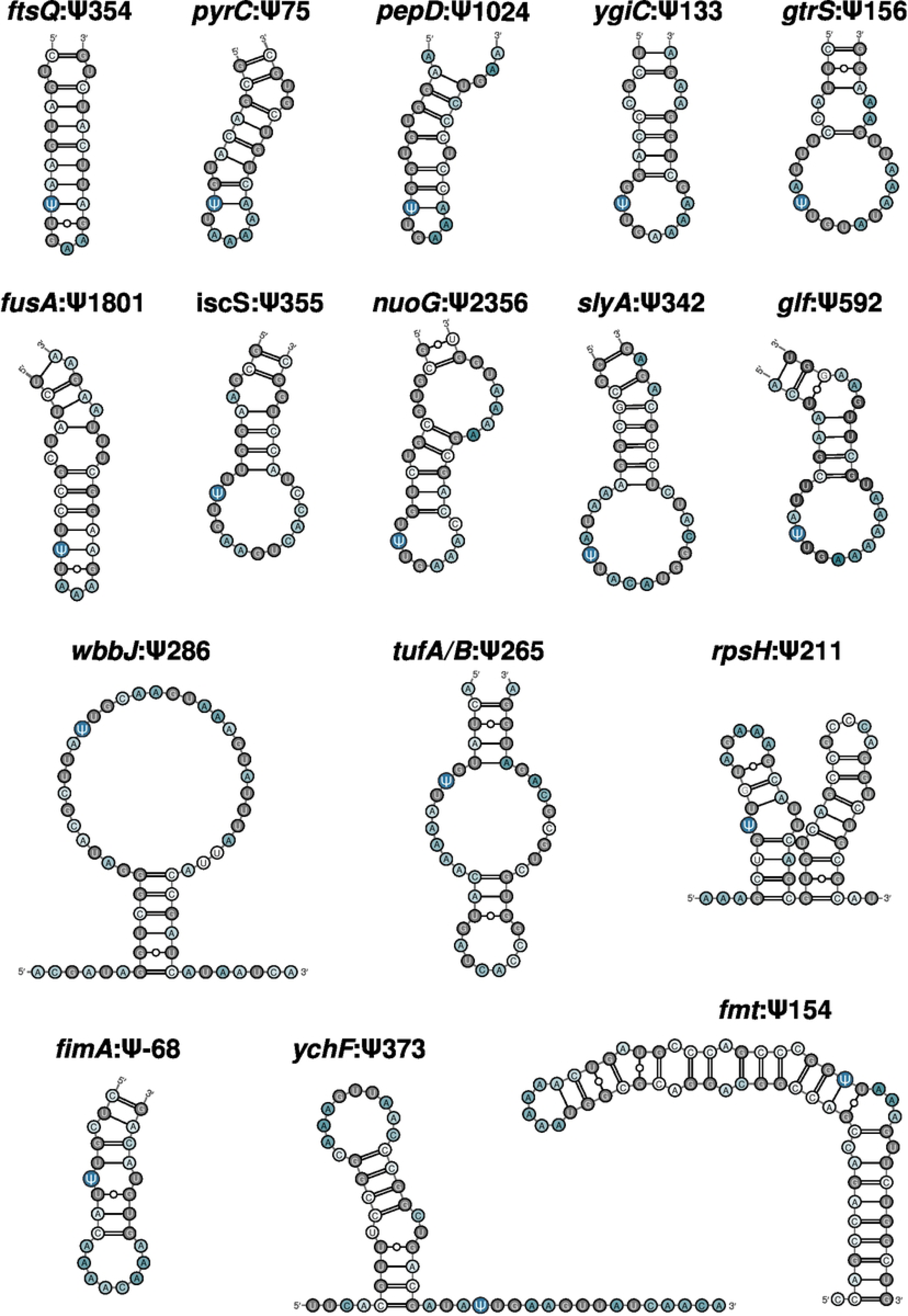
Predicted structures of additional RluA-dependent sites. DMS-informed secondary structure predictions for all RluA-dependent sites not shown in Figure 4, carried out in the same way. White-to-green gradient indicates DMS reactivity. Non-reactive nucleotides are in gray.

## Supplementary Information Captions

**Supplementary Table 1.** Peak heights across all libraries for sites meeting the coverage cutoff. Event annotations and enzyme assignments are also included.

**Supplementary Table 2.** Summary of events detected by simulating higher read coverage. For each event, peak heights are reported for the top depth where it was detected.

## Materials and Methods

### Strains

Pseudouridine synthase knockout strains were derived from the Keio collection (42). Each knockout was transduced into an MG1655 background using P1 phage transduction, as described in (43). Plate lysates were generated by growing each strain to mid-log phase in LB+0.1% glucose, adding CaCl2 to a final concentration of 5 mM, and mixing 3 mL of this culture with 50 μL of an existing P1*vir* stock. This mix was allowed to adsorb for 20 minutes, then mixed with 5 mL of melted top agar and poured onto 2 plates (LB with 0.1% glucose, 2.5 mM CaCl2, 1.5% agar). The plates were incubated overnight, then top agar was scraped into 50-mL tubes and vortexed with 5 drops of chloroform. To store the new P1*vir* stock, the tube was centrifuged for 10 minutes and the supernatant transferred to a new tube, to be stored at 4℃. To perform the P1 transduction, 1.5 mL of overnight cultures of *E. coli* MG1655 in LB were pelleted and resuspended in 0.75 mL P1 salts (10mM CaCl2, 5mM MgSO4). 100 μL of the resuspended cells were mixed with 1–100 μL of the P1 lysate, and left to adsorb at room temperature for 30 minutes. 1 mL of LB and 200 μL of 1M sodium citrate were added to the mixture, then the cells were grown for 1 hour at 37℃. The cells were pelleted and resuspended in 100 μL LB, then spread on LB plates supplemented with 50 ng/mL of kanamycin.

### Growth conditions

Overnight cultures were grown in 5 mL EZ Rich MOPS complete media at 37℃. Cultures were back-diluted into 15 mL pre-warmed MOPS complete media, to a target OD600 between 0.10 and 0.15, shaking in glass flasks at 220 rpm at 37℃.

### RNA extraction

RNA was extracted following the RNA snap protocol (44). Briefly, cultures were transferred to 15-mL tubes and pelleted by centrifugation at 3220 ×*g*, at 4℃. The media was decanted, then the tubes were spun down for an additional minute. The remaining media was aspirated and the pellets were flash frozen in liquid nitrogen. For RNA extraction, 200 μL of RNA extraction solution (consisting of 18 mM EDTA, 0.025% SDS, 1% 2-mercaptoethanol, 95% formamide) were added to each pellet, followed by vortexing for 20 seconds at top speed. The mixture was transferred to 1.5-mL tubes and incubated at 95℃ for 7 minutes in a ThermoMixer. To remove cell debris, tubes were centrifuged at >18000 ×*g* at 4℃ for 5 minutes. Extracted RNA was precipitated in ethanol by mixing 50 μL of the supernatant with 200 μL of 10 mM Tris (pH 7.0), 75 μL of 3M sodium acetate (ph 5.3) and 825 μL of ethanol.

### Pseudo-Seq library preparation

Pseudo-Seq protocol was based on (1,14), with some adjustments for handling of the shorter transcripts in *E. coli*. Total RNA was purified using the QIAGEN RNEasy Plus Mini Kit, which depletes RNAs shorter than 200 nt and removes genomic DNA. rRNA was depleted using either RiboZero or MicrobExpress kits. RNA was fragmented by adding RNA fragmentation buffer, incubating at 95℃ for 22 seconds, then quenched with stop solution. (RNA Fragmentation Reagents). CMC treatment was performed by incubating 40 μL RNA in 5 mM EDTA at 80℃ for 3 minutes, then adding 100 μL of 0.5 M CMC in BEU buffer (7 M urea, 4 mM EDTA, 50mM bicine, then incubating at 40℃ for 45 minutes, shaking at 1000 rpm. Mock treatment was performed by incubating adding 100 μL BEU without CMC before incubating. RNAs were precipitated in isopropanol and resuspended in 30 μL of pH 10.4 buffer, then incubated at 50℃ for 2 hours to reverse CMC adducts from guanidines and uridines. RNA fragments were precipitated, then run on a 10% TBE-Urea gel, and fragments 100–115 nt long were excised and purified. RNA 3′ ends were dephosphorylated with T4 PolyNucleotide Kinase, at 37℃ for 1 hour. Enzyme was inactivated at 75℃ for 10 minutes. 3′ adapters were ligated onto RNA fragments using T4 RNA ligase 2 truncated (20 μL reaction with ∼3 pmols input RNA, 25% PEG, 5uM Linker-1 [5′ App/CTGTAGGCACCATCAAT/3ddC]) for 2.5 hours at 25℃. Ligated RNAs were purified by size selection. Reverse transcription (RT) was carried out with SuperScriptIII: adapter-ligated RNAs were annealed to the RT primer (oCSB76, CAAGCAGAAGACGGCATACGAGATATTGATGGTGCCTACAG) in 11 μL of water with 25 pmol of the RT primer, incubated at 65℃ for 5 minutes. Reverse transcription proceeded at 50℃ for 30 minutes, with 0.5 mM dNTPs, 5 mM DTT, 10 U/uL SuperScriptIII and 1 U/uL SuperaseIn. NaOH was added to a final concentration of 0.1M, and the mixture was incubated at 95℃ for 20 minutes to hydrolyze template RNA. This solution was mixed with 20 μL of 2X RNA loading dye, and loaded onto a 10% TBE-Urea gel. Products 66-116 nt long, corresponding to insert sizes of 25-75 nt, were excised and purified. An adapter (oCSB77, /5Phos/NNNNNNNNNNAGATCGGAAGAGCGTCGTGTAGGGAAAGAGTGT/3SpC3/) was ligated to the 3′ ends of cDNAs as follows: cDNA was incubated for 2 minutes at 75℃ with 8pmols of the adapter and 1 μL of DMSO in a reaction volume of 10.8 μL. Then the rest of the ligation components were added (final reaction volume of 23 μL, 0.9 mM ATP, 20% PEG-8000, 1 μL T4 RNA ligase, high concentration), and the reaction proceeded overnight at 25℃. Partway through the incubation, the reactions were supplemented with an additional 0.5 μL of ligase. The reactions were precipitated, and ligated cDNAs were purified by size selection. After PCR amplification, libraries were sequenced on an Illumina NextSeq, generated 75-nt single-end libraries.

### REnd-Seq library preparation

REnd-Seq was performed as described in (45). rRNA was depleted from 10 ug total RNA with MICROBexpress, using two reaction volumes. Reaction was precipitated in isopropanol. RNA was fragmented using RNA Fragmentation reagents, by first incubating RNA in 40 μL water at 95℃ for 2 minutes, then adding Fragmentation Buffer to 1X and incubating for 25 seconds at 95℃ and finally quenching with Stop Buffer. Samples were precipitated in isopropanol and resuspended in 5 μL of 10 mM Tris 7. To size-select, RNA was mixed with one volume of 2X TBE-Urea sample loading buffer and run on a 15% TBE-Urea gel. RNA fragments 15-45 long were excised and purified, then precipitated. RNA was dephosphorylated with T4 PNK in a 20- μL reaction, incubated at 37℃ for 1 hour, then inactivated at 75℃ for 10 minutes. RNA was precipitated in isopropanol. Linker-1 was ligated to 3 moles RNA 3′ ends using T4 RNA ligase 2 (truncated), in a 20-μL reaction containing 25% PEG-8000, 1X T4 RNA ligase buffer, and 100 pmol of Linker-1. Reactions were precipitated in isopropanol, and resuspended in 5 μL 10 mM Tris pH 7. To purify ligated RNA, samples were mixed with one volume of 2X loading dye and ran on a 10% TBE-Urea gel. Ligated fragments were excised, purified, and precipitated. Reverse transcriptions were carried out with SuperScript III. First, samples were incubated with 25 pmol of RT primer (oCJ485) in 12 μL water at 65℃ for 5 minutes, then placed on ice. Remaining reaction reagents were added to the sample for a final volume of 20 μL (1X First Strand Buffer, 0.5 mM dNTPs, 5 mM DTT, 20U SuperaseIn and 1 μL SSIII), and incubated at 50℃ for 45 minutes. NaOH was added to a final concentration of 0.1M, and the mixture was incubated at 95℃ for 20 minutes to hydrolyze template RNA. This solution was mixed with 20 μL of 2X RNA loading dye, and loaded onto a 10% TBE-Urea gel. cDNAs were excised, purified, and precipitated.

### Pseudo-Seq data analysis

Adapter sequences were trimmed from read 3′ ends using bbduk (parameters: ktrim=r k=10 hdist=1 mink=1). 10-nt unique molecular identifiers (UMI) were removed from the 5′ end of the reads and added to the record headers using a custom script (parseBarcode.py). Reads were mapped to the *E. coli* MG1655 genome (NC_000913_3) using bbmap (slow=t perfectmode=true k=11 ambiguous=all). Alignments were converted to bam files and sorted with samtools, then filtered to keep only uniquely mapping reads. Multi-mapping reads were parsed into a separate bam file. PCR duplicates were collapsed by means of the UMI, using a custom script (dereplicate.py). Read 5′ ends were quantified on each strand using bedtools (bedtools genomecov −d −5-strand +), and were then saved as wig files.

### Peak calling

We calculate a Z-score for each U position relative to the 100-nt window surrounding it, requiring that at least 100 reads map to the window. Before the Z-score calculation, we modify the background window in two ways. First, we determine whether the window spans a strong transcript end. If it does, we shift the background window to avoid the peaks to achieve the following criteria: the 5′ boundary of a window must be at least 40-nucleotides downstream of the nearest transcript 5′ end, and at least 5 nucleotides downstream of the nearest transcript 3′ end. On the other side, the 3′ boundary of a window must be at least 25 nt upstream of the nearest downstream 3′ end, and at least 5 nt upstream of the nearest downstream transcript 5′ end. Transcript end positions were obtained from wild-type REnd-Seq data in the same growth conditions. After adjustment of the background window (or if adjustment was not necessary), we apply 95% winsorization to the window.

Z-scores are calculated relative to the adjusted 100-nt background window. To call CMC- dependent peaks, we require a Z-score ≥ 15 in the CMC-treated library, and for the CMC- treated Z-score to be at least 3-fold greater than the corresponding Z-score in the mock-treated library (Fig 2b). A CMC-dependent site is assigned as dependent on a given pseudouridine synthase if it meets three criteria: (1) it has sufficient coverage in the knockout library, (2) the knockout peak Z-score < 10, and (3) the WT CMC Z-score is at least 4-fold greated than the knockout peak.

### Structure analysis

In-cell DMS-Seq reactivities for wild-type *E. coli* were obtained from (33). Coordinates were converted to NC_000913_3 using LiftOver. For each site of interest, reactivities for the surrounding 100-nt window were parsed from this dataset into individual files (since only As and Cs react with DMS, other bases had a placeholder value of −999). These were provided as input to RNAfold for DMS-informed secondary structure prediction, using the --shape option (with parameters −p, −T 37, --shapeMethod=Z). Pairing probability matrices were parsed from the RNAfold output files, and columns were summed to summarize pairing probability. Pairing probabilities were visualized by calculating relative positions relative to the site of interest, and visualizing the pairing probability distributions for groups of sites at each relative position.

### Background site selection

A set of background sites was selected to control for expression level and for the influence of the RluA motif. First, we used our REnd-Seq data set to obtain WT expression levels for each gene that contains a high-confidence Ψ site. We consider only uridine positions within genes with RPKM ≥ 105 in our wild-type dataset, which was the lowest expression level among the genes containing high-confidence sites. Within these genes, we search for instances of the UURAA motif, and remove any sites that are within 100-nt of a Ψ site.

Secondary structure diagrams were generated using VARNA (v3.93)

### Soft-agar assay

Soft-agar plates, consisting of LB with 0.25% agar were poured the day before. Overnight cultures of target strains were diluted to OD600∼0.005 in 5 mL plain LB, shaking at 220 rpm at 37℃. Culture OD’s were monitored regularly, and plates were inoculated when OD600 reached 0.1. To inoculate, 1 μL of the culture was pipetted into the center of the plate. Plates were incubated at 37℃ for 8 hours, then plates were photographed and colony diameters were measured.

## Acknowledgements

We thank the members of the Gilbert lab and Li lab for helpful discussions; Tania Baker for sharing Keio collection strains. Sequencing was performed at the MIT BioMicroCenter, directed by Stuart Levine.

